# Zebrafish skeletal muscle cell cultures: Monolayer to three-dimensional tissue engineered collagen constructs

**DOI:** 10.1101/2020.12.10.419168

**Authors:** K.K Vishnolia, N.R.W Martin, D.J Player, E Spikings, M.P Lewis

## Abstract

Zebrafish (*Danio rerio*) are a commonly used model organism to study human muscular myopathies and dystrophies. To date, much of the work has been conducted *in vivo* due to limitations surrounding the consistent isolation and culture of zebrafish muscle progenitor cells (MPCs) *in vitro* and the lack of physiologically relevant models.

Here we report a robust, repeatable, and cost-effective protocol for the isolation and culture of zebrafish MPCs in conventional monolayer (2D) and have successfully transferred these cells to 3D culture in collagen based three-dimensional (3D) tissue-engineered constructs. Zebrafish MPC’s cultured in 2D were consistently reported to be Desmin positive reflecting their muscle specificity, with those demonstrating Desmin positivity in the 3D cultures. In addition, mRNA expression of muscle markers specific for proliferation, differentiation and maturation measured from both monolayer and 3D cultures at appropriate developmental stages were found consistent with previously published from other species *in vitro and in vivo* muscle data.

Collagen constructs seeded with zebrafish MPC’s were initially characterised for optimal seeding density, followed by macroscopic characterisation (three-fold contraction) of the matrix. Direct comparison between the morphological characteristics (proportion of cells) and gene expression profiles of cells cultured in collagen constructs revealed higher maturation and differentiation compared to monolayer cultures. In this regard, cells embedded in 3D collagen constructs revealed higher fusion index, Desmin positivity, hypertrophic growth, myotube maturity and myogenic mRNA expression when compared to in monolayer.

In conclusion, these methods and models developed herein will facilitate *in vitro* experiments, which would complement *in vivo* zebrafish studies used to investigate the basic developmental, myopathies and dystrophies in skeletal muscle cells.

## Introduction

The advantages of using zebrafish (*Danio rerio*) as an animal model for the study of human diseases are numerous. These include a high weekly reproduction rate, visualisation of organogenesis due to optical transparency and a large degree of genetic homology with humans^1-3^. As such, a vast number of disease models have been established in zebrafish, which accurately represent the phenotypes associated with human disease^4-10^.

A number of studies comparing mammalian and teleost systems provide evidence to support the use of zebrafish as a model to study the mechanisms of human myogenesis^11-13^, postnatal muscle function^14, 15^ and repair^16, 17^. Furthermore, it has been characterised that many of the same key signalling molecules are involved in these processes^1^. The work described here focuses on the isolation, characterisation, and culturing of zebrafish muscle precursor cells (MPCs), which are the progeny of satellite cells. These cells are skeletal muscle stem cells, which remain quiescent beneath the basal lamina of skeletal muscle fibres and play a vital role in repair, regeneration and growth of the tissue^18^. Upon injury, these cells proliferate extensively and undergo asymmetric cell division, which regulates the return to quiescence for a subset of cells, whilst a number of daughter cells terminally differentiate to form new muscle fibres or provide additional nuclei to existing fibres^19^. These mononuclear MPCs can be isolated through either tissue explant or enzymatic digestion and are routinely isolated from a number of mammals and teleosts e.g. salmon. Once isolated, MPCs can be subsequently cultured and used for *in vitro* experimentation, where cellular responses can be observed and manipulated in a highly controlled manner.

Norris et al. (2000) isolated blastomeres from zebrafish embryos (which in culture commit to form myoblasts) and studied the effect of Sonic Hedgehog signalling on the induction of a slow fibre fate, showing myotube formation after 48 hours in culture evidenced by myosin heavy chain expression^20^. Moreover, in 2011 Alexander and colleagues for the first time demonstrated the isolation MPCs from adult zebrafish and used microarray analysis to investigate gene expression over the time course of MPC differentiation to form myotubes^21^. The authors showed increased levels of the myogenic regulatory factor *myogenin* across a 14-day culture period, along with elevated mRNA levels of the muscle specific cytoskeletal protein, Desmin. These data suggest that MPCs can be successfully isolated, differentiated and matured over time in monolayer^21^. Whilst these data are informative, there remains a need to provide extensive morphological characterisation of zebrafish MPCs as to how they are differentiated *in vitro*, which can be used alongside transcriptional data to provide a basis for the future use of zebrafish muscle cells in research. This is of importance when seeking to use zebrafish skeletal muscle cells to investigate mechanisms that regulate regeneration, hypertrophy, and atrophy.

Furthermore, to-date MPCs isolated from fish species have been limited to conventional monolayer culture, which does not account for the three-dimensional (3D) nature of skeletal muscle *in-vivo*^*22*^. Indeed, many of the myopathies studied in zebrafish are diseases whereby the interaction between the sarcolemma membrane and the extra-cellular matrix (ECM) is disturbed^23^. Thus, it is important to develop a model system whereby isolated MPCs and the extra-cellular matrix can be manipulated, to investigate their independent and connected roles in muscular dystrophies.

To further develop conventional monolayer cellular systems to understand the pathophysiological mechanisms downstream, tissue engineered 3D skeletal muscle models have been developed. Such models aim to recreate the native *in vivo* structure of muscle tissue, *in vitro*. Our group employs a collagen-based model, which is now extensively published in the literature across multiple mammalian species^24-28^. These data suggest that constructs display highly aligned and differentiated myotubes from human^24^, rat and immortal mouse cell line sources (C_2_C_12_)^27, 29^. However, this model is yet to be tested with cells derived from non-mammalian species, providing scope for its use with isolated zebrafish MPCs.

The aim of the present work was to isolate and culture MPC’s derived from adult zebrafish and develop a collagen based 3D tissue engineered culture system that would allow these cells to form myotubes in a 3D formation that is more representative of their state *in vivo*. This information would allow for the routine utilisation of *in vitro* zebrafish muscle cell cultures and tissue engineered models, which could complement *in vivo* experimentation and shed further light on the cellular and molecular mechanisms underpinning skeletal muscle diseases in future.

## Methods

### Zebrafish maintenance

All the experiments were conducted as per the ethical animal welfare laws and guidelines from University of Bedfordshire, Luton, UK and were reviewed by the local ethical committee at the institute. Adult wildtype (AB-wildtype) zebrafish (10-12 months) were maintained in 40 litre (L) glass tanks at 28 LJC. Male and female fish were housed together in tanks and were fed three times a day with TetraMin^®^ (Tetra, Germany) flake food and once with freshly hatched brine shrimp (Artemia saline, ZM systems, UK).

### Isolation of skeletal muscle cells

Building on a previously published method for the isolation of skeletal muscle cells from salmon^30^, a protocol for the isolation of zebrafish skeletal MPCs was developed. The in-house protocol selected was as follows; 15-20 adult zebrafish (6–12 months old, mixed gender) were collected and washed twice with distilled water. Zebrafish were sacrificed with a lethal dose of MS-222 (Ethyl 3-aminobenzoate methanesulfonate) (Sigma-Aldrich, UK), followed by washing with 0.5% sodium hypo chloride or bleach (Sigma-Aldrich, UK) diluted in double-distilled water (ddH_2_O) for 45 seconds to remove any contaminates/oils from the surface of fish. Bleach residues were removed from anesthetised fish with three washes in phosphate-buffered saline (PBS). All internal organs and skin were removed on a sterile petri dish using a sterile scalpel. Further the soft muscle tissue (5 grams per isolation) was moved to a biological safety cabinet and washed twice with PBS. Tissue was minced with sterile scalpels in a petri dish and subsequently incubated in a sterile 50 ml tube with 10 ml of 5 mg/ml collagenase type 1A solution (Sigma-Aldrich, UK) at room temperature for 45 minutes on a shaker at 200 revolutions per minute (rpm). 20 ml of isolation media i.e. L15 medium supplemented with 0.8 mM calcium chloride (CaCl_2_), 2mM glutamine, 3 % fetal bovine serum (FBS), 100 µg/ml penicillin/ streptomycin (Sigma-Aldrich, UK) was added after incubation in order to stop the collagenase activity. Subsequently, the cell suspension was filtered through 100 and 40 µm nylon strainers (Fisher Scientific, UK). The filtrate was centrifuged at 1000 x g for 10 minutes and the supernatant was discarded. The pellet was re-suspended in 4 ml PBS prior to layering over 4 ml of Ficoll solution (GE Healthcare life sciences, UK) in a 15 ml centrifuge tube and centrifuged at 1400 x g for 45 minutes. Mononuclear cells were collected from the top of Ficoll layer and washed with PBS followed by re-suspension in 10 ml growth media i.e. L15 supplemented with 0.8 mM CaCl_2_, 2mM glutamine, 20 % FBS, 100 µg/ml penicillin/ streptomycin. Cells were counted using a haemocytometer and plated on coverslips pre-coated with 0.2% gelatin in six well plates. These were then cultured at 28 °C in a sterile tissue culture incubator.

Once the cells reached 90% confluency (approximately in three to four days), cells were pushed for differentiation to fuse and form multinucleated myotubes using low serum differentiation media i.e. L15 supplemented with 0.8 mM CaCl_2_, 2mM glutamine, 2 % horse serum, 100 µg/ml penicillin/ streptomycin.

Morphological characterisation in monolayer was conducted on early formed myotube i.e. after 3-4 days in differentiation medium (time point denoted as “early myotubes”) and late-stage when ideally myotubes should be matured by aggregating more nuclei per myotube i.e. after 12-14 days in differentiation medium (time point denoted as “myotubes”). Morphological characteristics, fusion efficiency and Desmin positivity were computed from 20 different isolations and six randomly selected Desmin (for Desmin protein) and dapi (for nuclei) immune stained images per isolation.

### Tissue engineered collagen constructs

The collagen constructs used in the present study are based on previously published protocols from our group^24, 25, 27^. Briefly, 3D collagen gels were formed by seeding isolated zebrafish MPCs at the required plating density (see results section) into a 1.5 ml collagen-10 x Minimal Essential Media (MEM) solution (collagen: First Link, Birmingham, UK; Minimal Essential Media [MEM], Fisher, UK) neutralised by sodium hydroxide (NaOH). 1.5 ml total volume of collagen gel was divided in 1.3 ml collagen, 150 µl of the MEM solution, followed with addition of cells at required density in 50 µl volume. The neutralised solution was allowed to polymerise within a chamber between two fixed points for 15 minutes at 28°C. The two fixed points originated from a custom-made structure, termed floatation bars and A-frame. Floatation bars were constructed from small pieces (6 mm x 9 mm) of polyethylene plastic mesh (Darice Inc, Strongsville, Ohio, US) tied with 0.3 mm diameter stainless-steel wire (Scientific Wire Company, Great Dunmow, UK). A-frame was constructed by bending a 0.7 mm stainless steel wire (Scientific Wire Company, Great Dunmow, UK) and fixing it in between the floatation bar. Sylgard barriers were used to divide the glass chamber and keep the collagen gels in confined space (20mm x 12mm x 10mm) (Figure 1). Flotation bars and A-frames, along with the glass chambers and Sylgard barriers were sterilised before experimentation in 70% ethanol and UV light in a biological safety cabinet for 30 minutes, respectively.

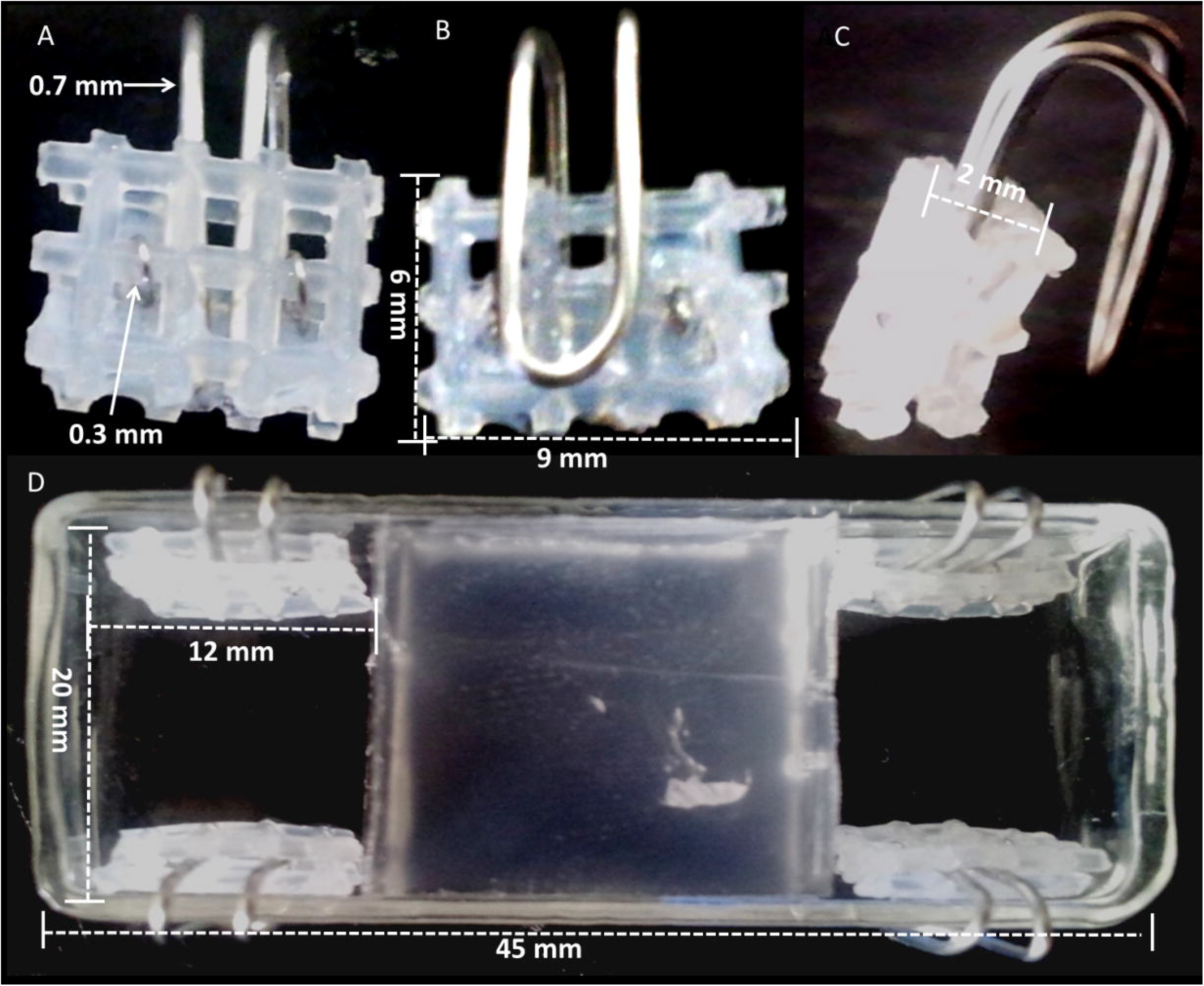

#### Immunohistochemistry

##### Cells in monolayer

Media was aspirated from the culture wells followed by two PBS washes. Cells were fixed for 10 minutes in 50% PBS and 50% ice cold 1:1 methanol-acetone (Fisher Scientific, UK) solution and then exposed to 100% 1:1 ice-cold methanol-acetone solution for a further 5 minutes. Subsequently, coverslips were removed from the wells and cells were treated for 45 minutes with 100 µl of blocking solution prepared using Tris Buffered Saline (TBS, pH 8.5), 5% goat serum (Sigma-Aldrich, UK) and 0.2% Triton-X 100 (Fisher Scientific, UK). Cells were then rinsed with TBS prior to overnight incubation with 100 µl of rabbit polyclonal, anti-Desmin (Ab86083, Abcam, UK) primary antibody at a dilution of 1 in 200 in TBS, 2% goat serum and 0.2% Triton-X 100. The following day after washing the coverslips three times with TBS, they were incubated in goat anti-rabbit IgG, TRITC (Abcam, UK) secondary antibody diluted at 1 in 200 with TBS, 2% goat serum and 0.2% Triton-X 100 for one hour, followed by three washes with PBS. 4’, 6-diamidino-2-phenylindole (DAPI) (Sigma-Aldrich, UK) was used to stain nuclei, diluted at 1 in 3000 in deionised water and incubated for 10 minutes, followed by three washes in PBS. Coverslips were finally mounted onto glass microscopic slides (Fisher Scientific, UK) using a drop of MOWIOL (Sigma-Aldrich, UK) mounting medium containing the anti-fade agent DABCO (Sigma-Aldrich, UK). The coverslips were viewed and imaged using an inverted laser scanning confocal microscope (Leica, UK).

##### Tissue engineered 3D collagen constructs

Collagen tissue engineered constructs were fixed *in situ* like monolayer coverslips, but with an increased incubation time (20 minutes). Following fixation, gels were detached from their fixed points and mounted on poly-l-lysine coated microscope slides (Fisher Scientific, UK) and the constructs were ringed with PAP pen (Fisher Scientific, UK). Gels were whole mount stained following the same protocol used for monolayer cultures with the same antibodies. Constructs were blocked for three hours in the blocking solution, before being washed three times using TBS. The anti-Desmin primary antibody solution at 1 in 200 dilution, in a volume enough to submerge the gels, was pipetted on the gels and incubated overnight in a humidified chamber at room temperature to avoid evaporation of the antibody and drying of the constructs. Next day, primary antibody was removed, and the constructs were washed three times in TBS before incubating the gels for three hours in the goat anti-rabbit IgG, TRITC secondary antibody solution at 1 in 200 dilution. After secondary antibody incubation, the constructs were washed three times with PBS prior to the addition of DAPI for 30 minutes. Finally, constructs were washed three times in PBS before mounting with glass coverslips (Fisher Scientific, UK) using MOWIOL containing DABCO. Immuno-stained engineered constructs were imaged using an inverted confocal microscope (Leica, UK).

### Gene expression analysis

#### RNA extraction and PCR

RNA was extracted at different time points throughout experiments using the TRIzol (Life technologies, UK) method, following the manufacturer’s instructions. Cells in monolayer cultures were scraped and homogenised, whereas collagen constructs were homogenised using a hand-held stick homogeniser (IKA T10 Fisher Scientific, UK) after being washed several times in PBS. RNA (1 µg) was transcribed using the Precision mqScript Reverse Transcription Kit (Primerdesign Ltd, UK) according to the manufacturer’s protocol and the cDNA was diluted 1:2 in molecular biology grade water (Sigma,UK). Gene expression analysis was performed by quantitative real time PCR on a RotorGene 6000 cycler (Corbett Research, UK) using a 72 well rotor. Gene expression data was generated from three different isolation cultures with three technical replicates for each isolation in monolayer or 3D collagen constructs. Relative gene expression data for each muscle marker was normalised against two house-keeping genes *ef1*α and β*-actin* (Primer sequences are detailed in Table 1). All results are presented relative to a single designated control sample labelled as ‘after isolation’ using the Rotor Gene software (Version 1.7, Corbett research), as previously published by our group^31^.

Expression of different myogenic markers were measured at different stages of development and differentiation from both monolayer (time points labelled as sub-confluent, confluent, early myotubes and myotubes) and 3D tissue engineered collagen constructs (time points labelled as before differentiation and myotube stage). Expression of myogenic regulatory factors *myoD, myogenin, myf6*, along with myogenic inhibitor *myostatin*, hypertrophic gene insulin like growth factor-1 (*igf-1*), slow (*smyhc1*) and fast isoform of myosin heavy chain (*myhc4*), were investigated.

#### Statistical analysis

A one sample Kolmogorov-Smirnov statistical test was used to determine the normality of data from all the replicates (where necessary after logarithmic [log_10_] transformation). One-way ANOVA with Bonferroni post-hoc correction was used to determine any statistical differences among time points and conditions investigated. Data are expressed as mean ± standard deviation, with significance set at p< 0.05.

## Results

### Monolayer culture: Isolation and culture conditions of zebrafish MPCs

Enzymatic digestion of 5 grams muscle tissue from 15-20 adult zebrafish (both male and female) in 5 mg/ml type 1 collagenase resulted in an average cell yield of 14.8 × 10^6^ ± 1.4 × 10^6^ cells. Approximately 300,000 cells/well were plated in six well plates. Increase or replication of cells can be evidently observed from the immuno-stained images in Figure 2 where cells are scattered at day 1 (Fig 2-A), whereas they are almost 90% confluent at day 4 (Fig 2-B). Myogenic purity or Desmin positivity of this cell population was determined to be 42 ± 7.80% at day 1. Early myotubes (2-3 nuclei) were observed after 3-4 days of switching to differentiation media (Figure 2-C) confirming the fusion capacity of isolated cells. Matured myotubes with increased number of nuclei per/myotube were observed after 12-14 days in differentiation media (Figure 2-D). Of note, we were successfully able to passage and/or freeze-thaw and sub-culture the zebrafish MPC’s, however all the data included in this study was generated from primary cell cultures.

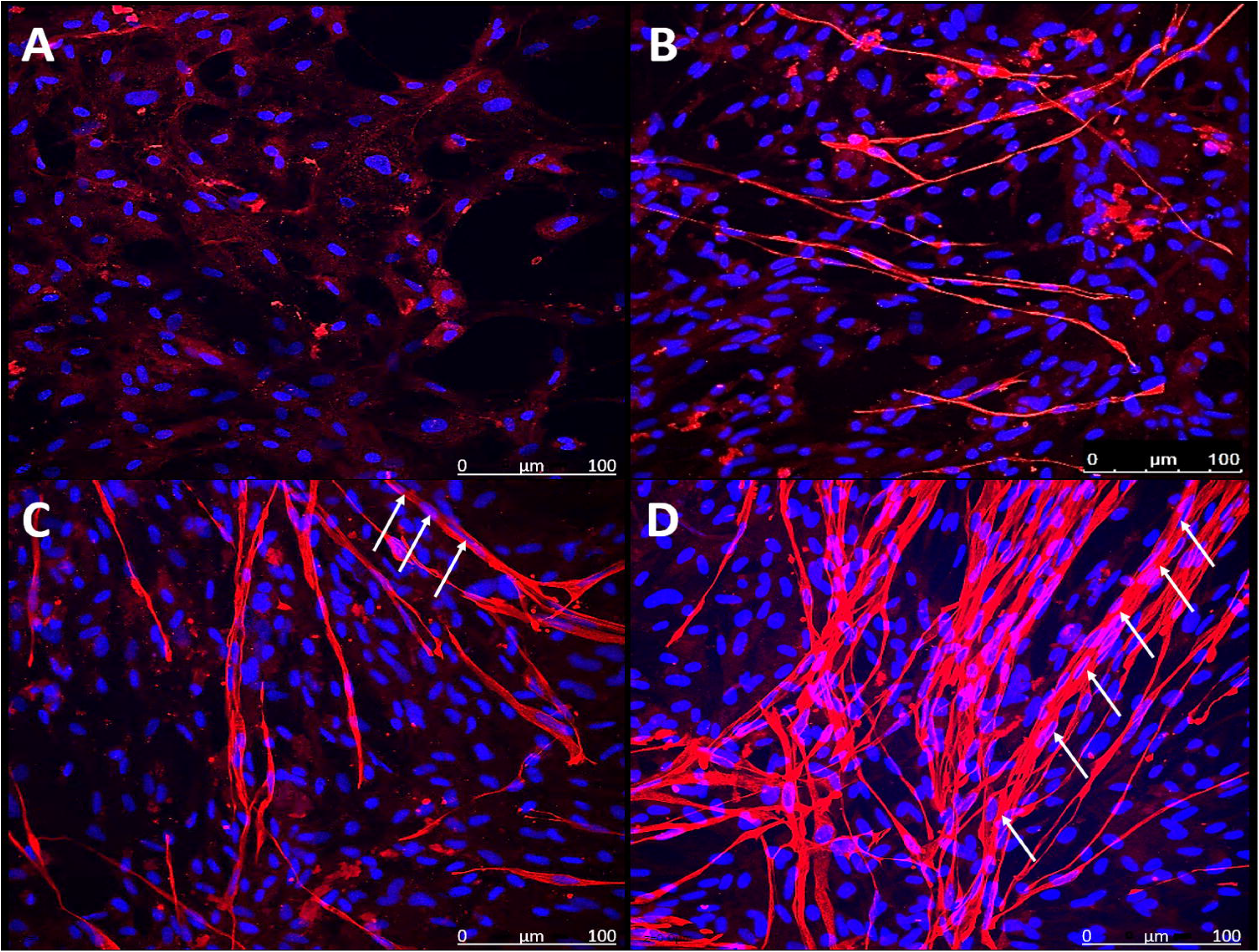

To further analyse differentiation of zebrafish MPCs at the molecular level, we measured the mRNA expression of the muscle regulatory factors *myoD, myogenin* and *myf6. myoD* mRNA levels were significantly increased in confluent cells (3-4 days after plating) compared to sub-confluent cultures (1 day after plating) (1.62 ± 0.18 vs 1.16 ± 0.17, p < 0.05), and were further augmented in early myotubes (3-4 days in differentiation media) compared to all time points (4.85 ± 0.56, p < 0.0001), before attenuating towards basal levels in myotubes stage (12-14 days in differentiation media) (1.61 ± 0.30) (Figure 3). A similar pattern was observed for *myogenin* expression, which significantly increased as cells transitioned from sub-confluence (0.07 ± 0.01; p < 0.05) to confluence (0.18 ± 0.02; p < 0.05) respectively, then to early myotubes (3.02 ± 0.32; p < 0.0001) and then reduced in later myotubes (0.78 ± 0.12; p < 0.0001) (Figure 3). *myf6* expression increased more dramatically from sub-confluence (0.62 ± 0.08) to confluence (1.94 ± 0.21, p < 0.05) before further augmenting in early myotubes (5.44 ± 0.44, p < 0.0001) and remaining elevated in myotubes (4.38 ± 0.75) (Figure 3). These results confirm that the zebrafish MPCs were committing to the myogenic lineage and undergoing fusion to form myotubes. Furthermore, these data support the roles of the measured myogenic regulatory factors in the specification of MPCs in early and late MPC differentiation in the zebrafish.

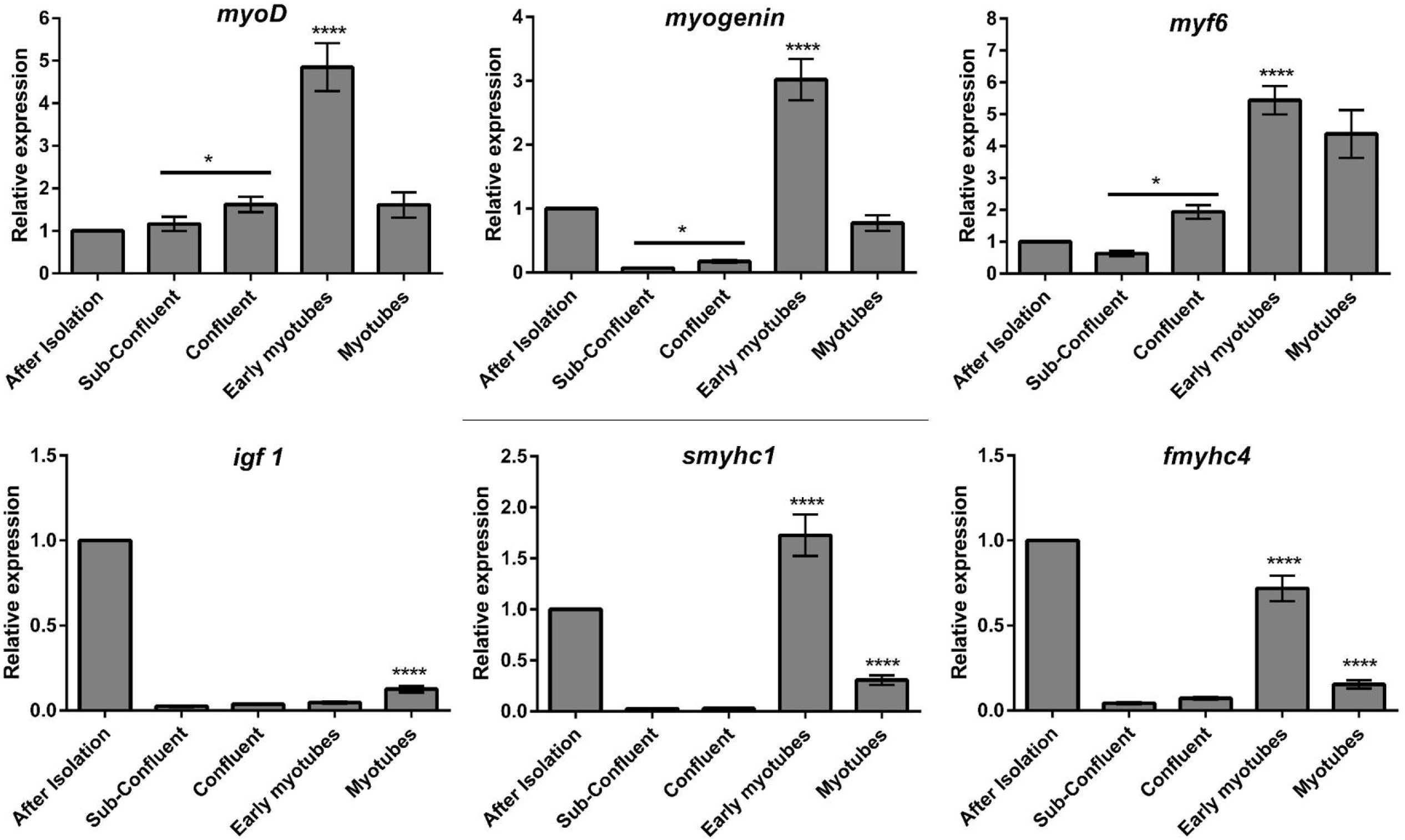

### Morphological characterisation of myotubes in monolayer

At early and late time points during MPC differentiation, myotubes were fixed and stained for Desmin protein and nuclei, to visualise cellular morphology (Figure 2A-D). Desmin positivity of a culture does not necessarily define the fusion capability of muscle cells in culture therefore fusion efficiency per isolation was quantified. Fusion efficiency was measured as a ratio between number of nuclei incorporated in myotubes and total number of Desmin positive nuclei. The overall fusion efficiency of the myogenic cells revealed that approximately 60% of myogenic positive cells in the cultures were able to form myotubes (Table 2), which remained consistent over time in culture. Similarly, the number of myotubes per image did not significantly change with time in differentiated cultures, was approximately six myotubes per image (Table 2). Interestingly, whilst overall levels of fusion did not change, both the length (360.92 ± 7.55 vs 336.87 ± 7.01μm) and width (10.80 ± 0.27 vs 8.30 ± 0.21μm) of myotubes were significantly increased in later myotubes versus early myotubes (p<0.05, Table 2). This suggests a degree of maturation and/or muscle growth during this period, which is mediated through increased protein accretion independent of myonuclear addition. To confirm this hypothesis, we generated a frequency distribution of myotubes containing various myonuclear densities at early and late myotube stages (Figure 4). Indeed, whilst there exists a small tendency for greater frequency of myotubes with increased nuclei in late myotubes, there was no significant interaction effect (p=0.415), therefore confirming that increases in myonuclear number are not significant in late versus early myotubes.

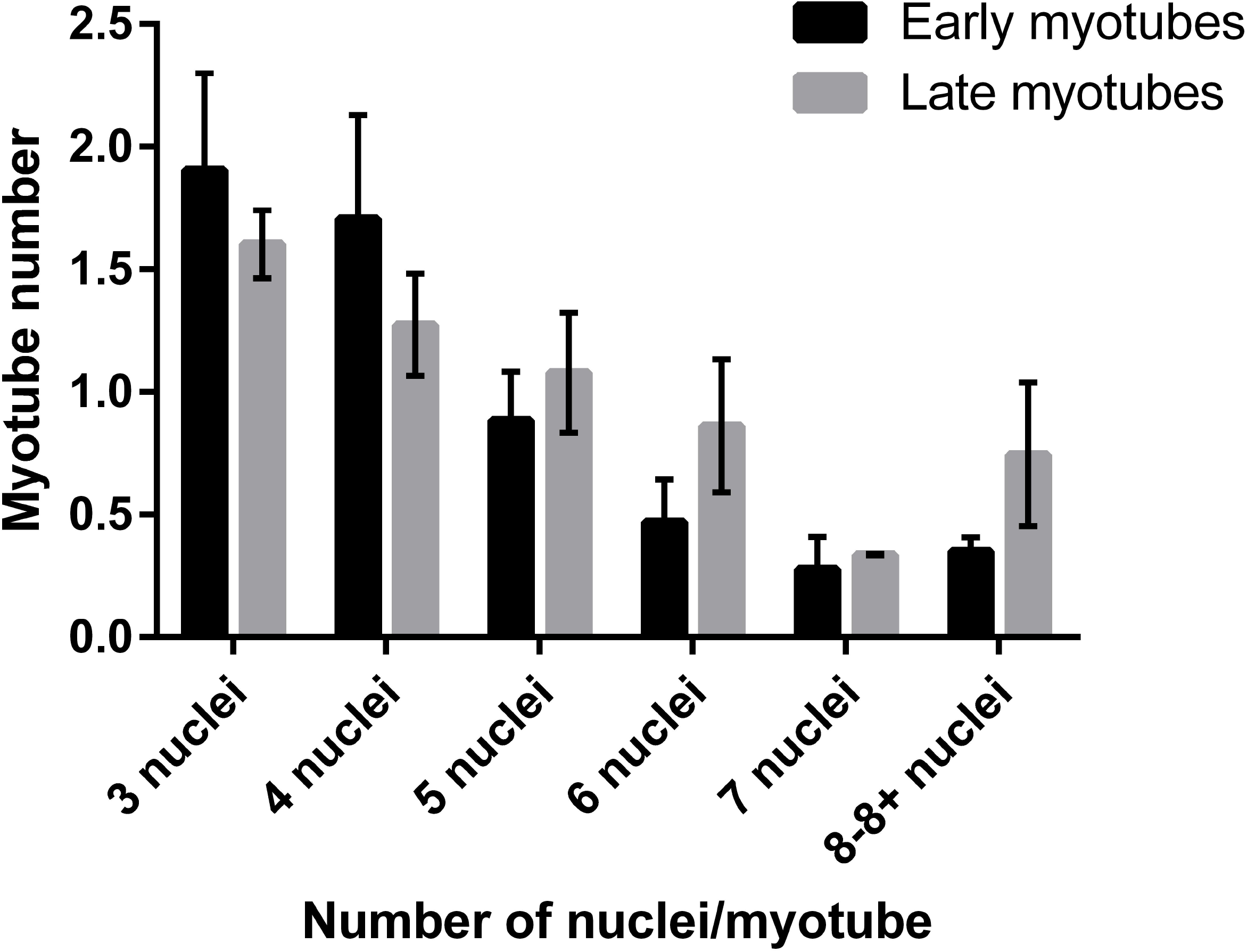

### Myosin Heavy Chains (MyHC) mRNA expression in monolayer

Myosin heavy chains form the thick filament within the contractile apparatus in skeletal muscle, where the expression of MyHC isoforms is different throughout the stages of development and maturation. We measured the expression of both, fast (*fmyhc4*) and slow myosin heavy chain (s*myhc1*) mRNA during the culture of zebrafish MPCs *in vitro. fmyhc4* expression did not increase significantly in cells from sub-confluence (0.04 ± 0.006) to confluence (0.072 ± 0.008, p > 0.99) but reached its peak expression in early myotubes (0.72 ± 0.08, with p < 0.0001) before levels attenuated in late myotubes (0.15 ± 0.024, p < 0.0001) (Figure 3). *smyh*c1 remained expressed at low levels in sub-confluent (0.025 ± 0.002) and confluent cells (0.028 ± 0.004) with no significant difference before increasing dramatically in early myotubes (1.72 ± 0.20, p < 0.0001) and thereafter reducing (0.30 ± 0.04, p < 0.0001) (Figure 3).

### Tissue engineered constructs: Zebrafish MPC morphology in 3D

Following characterisation in monolayer, freshly isolated zebrafish MPCs were seeded into 3D collagen constructs. Seeding density was optimised by plating the collagen gels at a range of seeding densities i.e. 4, 6, 8, 10 and 12 million cells/ml. Few cells were observed in phase contrast images of collagen gels seeded at 4, 6 and 8 million cells/ml, suggesting not enough adherent cells (Supplementary figure 1). Collagen gels seeded with 12 million cells/ml were found to be detaching from the anchor points or A-frames after first or second day in culture (data not shown). However, plating density of 10 million cells/ml showed optimum results in terms of maximum number of cells observed in phase contrast images and later confirmed with Desmin staining. Like monolayer, collagen gels seeded with zebrafish MPC’s were initially cultured in growth media for three to four days, before being switched to differentiation media after confirming their confluency.

Collagen gels seeded with 10 million cells/ml evidently demonstrated characteristic contraction in the width (gel bowing or contracting from the sides) over time as shown in Figure 5A-B. This phenomenon has already been reported in literature using different cell types and formats of collagen hydrogels^18, 24, 27, 32^. Contraction of collagen matrix width embedded with zebrafish MPCs was macroscopically documented over the course of 12 days i.e. from day 1 (Figure 5-A) till day 12 after differentiation (Figure 5-B) time points.

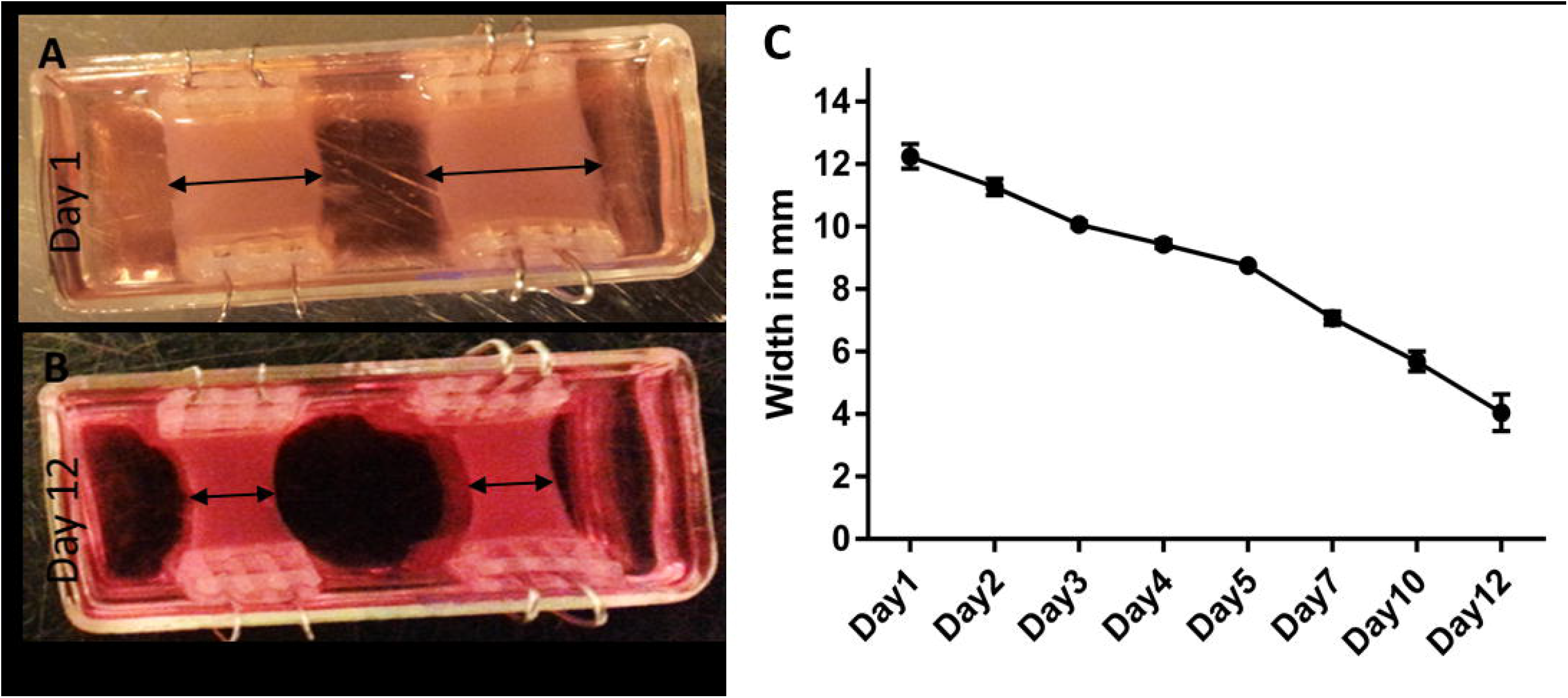

Tissue engineered collagen gels seeded with zebrafish MPC’s stained for Desmin before differentiation (day 1 in culture), revealed only single or mononuclear cells (Figure 6-A). In contrast, post differentiation immunostaining demonstrates multinucleated, unidirectional bundles of myotubes, as confirmed in Figure 6-B. The number of myotubes per microscopic frame from Desmin stained 3D collagen constructs (20 images/construct, n=6 collagen constructs), showed a significant increase from 1.55 (± 0.40) at differentiation stage to 19.11 (± 0.50) at 5 days post differentiation (p < 0.05, Figure 6C). Significantly higher number of myotubes were found aggregated with 5, 6, 7 and 8-8+ nuclei per myotubes at 5 days post differentiation time point, suggesting maturation of myotubes in the collagen constructs (p < 0.05, Figure 6D). Also, significant increment of fusion efficiency was observed in the collagen constructs i.e. 82% compare to monolayer i.e. 60% (p < 0.05, Figure 6E).

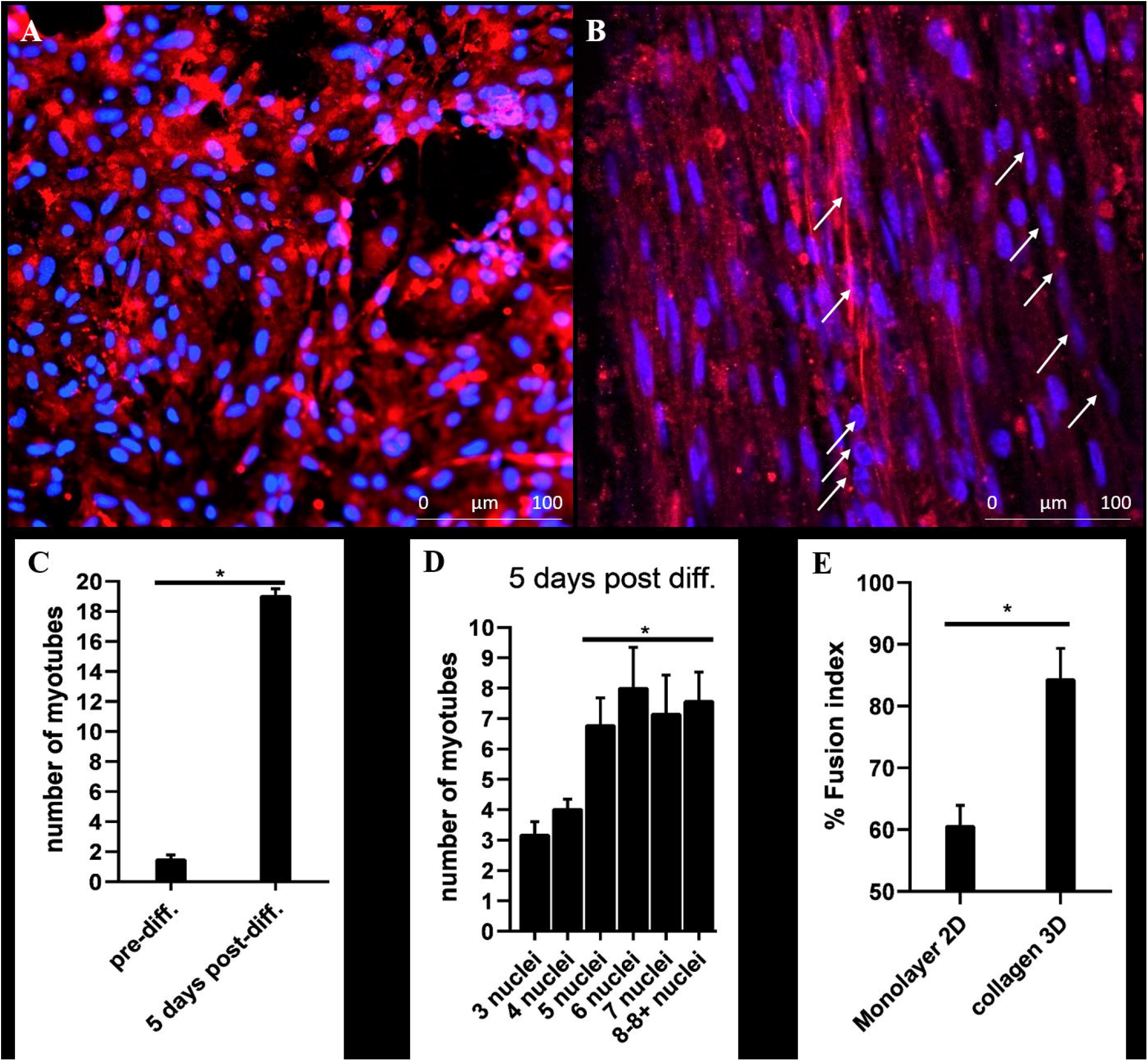

### Gene expression analysis in 3D collagen constructs

Gene expression analysis was conducted using a sample of MPCs just after isolation, as an external control. The greater number of myotubes present at the post differentiation time point was underpinned by a greater expression of myogenic genes. There was a significant increase in the expression of *myoD, myogenin*, and *myf6* following differentiation compared to pre-differentiation (p< 0.05, Figure 7), demonstrating the activation of these genes required for the differentiation and maturation process. When comparing the myosin heavy chain mRNA response in 3D compared to monolayer culture, only *smyhc1* showed a significant increase following differentiation (p< 0.05, Figure 7), while no difference was observed for *fmyhc4* (p>0.05, Figure 7). This suggests the myosin heavy chain response is specific and dependent on the culture type.

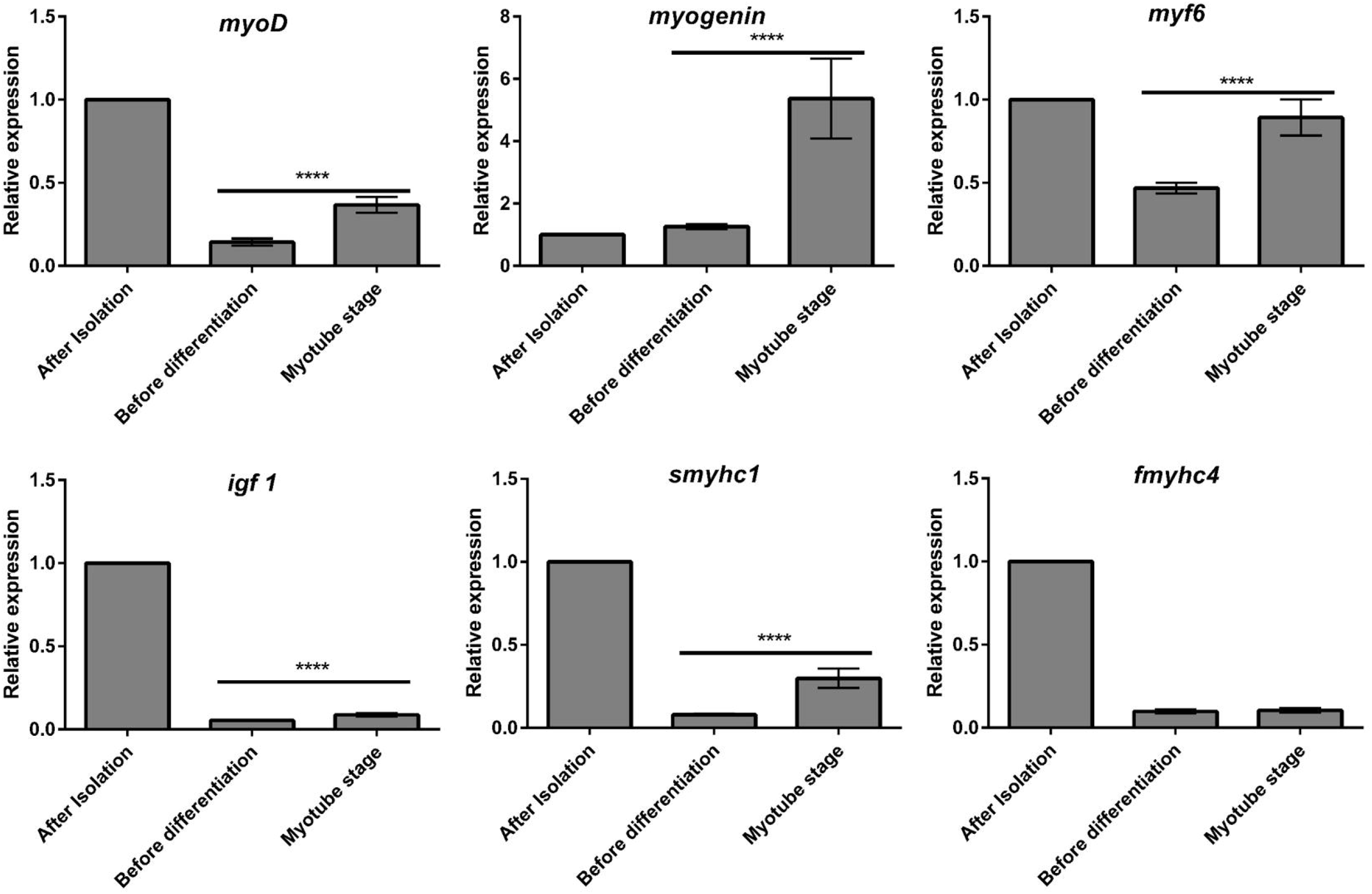

## Discussion

Although the protocol for the isolation of zebrafish skeletal muscle progenitor cells from muscle tissue has been reported by Alexander et al, 2011^21^, further characterisation of the morphology and myogenic gene expression is required. Furthermore, we sought to establish a zebrafish 3D culture model based on previous work of the group in cells derived from other species.

In the present work, we isolated a yield of approximately 14 million cells per isolation from 5 grams soft muscle tissue (15-20 adult zebrafish). Isolated cells warranted characterisation to ascertain whether myogenic cells were contained within the heterogeneous population. Approximately 42 ± 7.80% of the isolated cells were found to be Desmin positive, indicating myogenic potential in culture. The initial proportion is not as high as with other published protocols for different species, but both the culturing system in monolayer and 3D constructs allowed for replication and differentiation of these cells into myotubes in line with published literature from our group from different species and cell lines^29^. Primary cultures of MPCs typically result in varying levels of myogenic purity owing to several variables such as age, duration of enzymatic digestion, extent of tissue trituration, cell straining conditions to remove debris and centrifugation speed^32, 33^. Previously reported zebrafish cultures reported *myoD* mRNA expression levels of approximately 75% in isolated cells^21^, whilst human MPCs can range from 15-85% Desmin positivity^32, 34^. The isolated cellular population using the current protocol does not differentiate between cell types, therefore a heterogeneous population is isolated. Other than myogenic cells, it is likely that most of the non-myogenic population consists of lineages originating from the extracellular matrix. Future experimentation should seek to investigate whether different fractions of cell types (i.e. myogenic vs. non-myogenic^32^), would influence myotube formation.

Pure myogenic fraction of cells per isolation can be obtained after replicating and passaging them in monolayer and implementing various different techniques reported in the literature such as flow cytometry^35^, cell sorting, magnetic cell sorting^32^, differential cell plating^36^ and fractionation using Percol/ Ficol density gradient^33^. In conventional monolayer cell culturing, cells are already removed from their physical *in vivo* environment, therefore it becomes of increasing significance to move towards more biomimetic models for tissue culturing^32^.

Seeding densities play a crucial role in monolayer as well as three-dimensional tissue engineered constructs to allow sufficient cell-cell interaction, which is required for fusion^34^. Here, we observed that 1.5 ml collagen constructs seeded with 2, 4, 6, 8 million cells/ml did not adhere with the surrounding collagen, whilst constructs seeded with 12 million cells/ml detached from anchor points. However, cells seeded with 10 million cells/ml showed significant contraction of the gels over time and most importantly presence of straight, multi-nucleated bundles of aligned myotubes (Figure 6). Previous findings from our group using human MPCs at different plating densities in an acute cyto-mechanical model (culture force monitor), demonstrate the difference in cell-cell and cell-matrix interactions at different seeding densities^34^. The differences in cell-cell and cell-matrix interactions, may offer an explanation for the differences observed at different seeding densities presented here.

In zebrafish, bundles of multinucleated muscle fibres are arranged in parallel allowing movement and to sustain multimodal swim due to their contractile properties^13^. In 3D collagen constructs zebrafish MPC’s are provided with similar *in vivo* niche held between two fixed points. Isometric tension is formed between the fixed points due to contraction of cells as they attach to the matrix leading to generate mechanical stimuli facilitating reorganisation and alignment of myoblasts along lines of principle strain^24^. To further characterise and understand the molecular signals driving the differentiation of zebrafish MPCs, mRNA levels of the muscle regulatory factors *myoD, myogenin* and *myf6* were determined in isolated MPCs as they proliferated and underwent fusion. We showed a clear increase in the expression of all myogenic regulatory genes when differentiation was initiated in both monolayer and 3D, and a subsequent reduction in *myoD* and *myogenin* mRNA levels as myotubes matured in monolayer, whereas *myf6* expression remained elevated. These data match with those previously reported across species, showing that *myoD* is expressed in proliferating MPCs and is required for myogenic determination, and thereafter *myogenin* is important for terminal differentiation to form myotubes^37^. Furthermore, *myf6* levels which in the present work remained elevated in monolayer once fusion had occurred, have previously been shown to be expressed highly in post-regenerative muscle^38^.

Immuno-cytochemical analysis of the developed myotubes in early and late phases of monolayer culture clearly indicated that whilst no further cellular fusion occurred over time in culture, there were increases in myotube size (length and width). Further analysis of myonuclear quantity showed that nuclear accretion did not occur in myotubes over time. Therefore, this would suggest that increased myotube area was a product of growth, independent of cellular fusion. Increased growth of muscle fibres or myotubes can indeed occur without the need for myonuclear addition, through a net increase in protein synthesis^39^, however it is likely that extreme increases in muscle size will require new myonuclei in order to maintain the levels of transcription required for additional growth^39-41^. Approximately 60% fusion index in monolayer and 82% in 3D collagen hydrogels, suggests that a substantial number of myogenic cells remain un-differentiated in monolayer cultures. In contrast, over 82% of myogenic cells were able to differentiate and form multi nucleated myotubes, with much higher numbers of nuclei per myotube when cultured in more bio-mimetic 3D collagen constructs. Overall, improvement in hypertrophic growth, fusion efficiency, and alignment of zebrafish MPC’s in the physiological environment provided by 3D collagen constructs proves significant advancement over conventional monolayer culture system. With the stark differences in fusion and growth between the two models, it will be possible to investigate the physiological proportions of other cell types that contribute to ECM maintenance and the underlying functional mechanisms which prevail in zebrafish models of dystrophy.

Finally, we measured mRNA levels of the fast and slow myosin heavy chain isoforms throughout MPC differentiation in both monolayer and 3D, as an indication of levels of myotube maturation. We found the expression of both isoforms increased during myotube differentiation in monolayer, whilst only *smyhc1* increased in 3D following differentiation, suggesting a predominance of this myosin heavy chain exclusive to the 3D environment. The increase in expression of the *smyhc* in the collagen model, despite no increase in *fmyhc*, was in line with previous literature from our group, suggesting the myosin heavy chain expression may be dependent on the fixed 3D nature of the collagen^34^. It has previously been shown that during zebrafish myogenesis *in vivo*, embryonic slow muscle fibres co-express fast and slow myosin heavy chain isoforms before reaching full maturity^14^, and therefore the expression of both isoforms in monolayer is not unexpected. Furthermore, MPCs were isolated from a pooled muscle homogenate rather than a specific muscle fibre of slow/fast type, and therefore would likely contain a mixture of fibre types with associated MPCs, contributing to the expression of both fast and slow myosin heavy chains.

In conclusion, our study confirms that MPCs can be successfully isolated from zebrafish with myogenic capacity, replicating, growing, and differentiating *in vitro*. Moreover, these cells are capable of incorporation into a 3D tissue engineered collagen-based culture system where they have been shown to differentiate more in line with expectations based on markers expressed *in vivo* than their 2D cultured counterparts. The characterisation and establishment of zebrafish skeletal muscle 3D culture system will facilitate to complement several *in vivo* experiments conducted on zebrafish investigating skeletal muscle development, understand functional mechanisms, high throughput drug screening and disease modelling.

## Acknowledgements

We would like to acknowledge all the support provided by Dr. Jonathan McDermid, University of Leicester, UK for helping us with keeping and maintenance of zebrafish. We would also like to acknowledge Prof. Vivek Mudera from UCL, and our colleagues from Loughborough University for the critical discussions of project.

## Author Contributions

KV conceived, designed, performed, collected data, and wrote the manuscript. DP, NM contributed to critical analysis of data, manuscript revisions. ES, ML supervised the project, critically analysed the data, reviewed the manuscript, and helped in designing initial concept of project.

