## Supplementary material for "Zebrafish skeletal muscle cell cultures: Monolayer to three-dimensional tissue engineered collagen constructs": Figure legends

**Figure 1:** Floatation bars and A-Frame set up for 0.75 ml collagen gels. **A, B** and **C** represents the front, back and side view of custom-made floatation bar with A-Frame. **D**) Whole set up for two 1.5 ml collagen gels in a glass chamber separated by Sylgard spacer. Dotted lines represent the dimensions of A-Frames, whole chamber and the gels. Solid arrows annotate grade of wires used for making A-Frames.

**Figure 2:** Representative immuno-stained images of zebrafish skeletal muscle cells in monolayer at four different stages. Red represents Desmin protein stained by Desmin antibody and blue represents nuclei stained by DAPI. **A**) Zebrafish skeletal muscle cells at sub-confluent stage, **B**) Cells at confluent stage, **C**) Cells at early myotube stage, **D**) Cells at late myotube stage. Arrows in **C** and **D** highlights multi nucleated myotubes. Images were captured at 100X magnification.

**Figure 3:** Relative mRNA expression graphs of zebrafish skeletal muscle cells cultured in monolayer sampled at after isolation, sub-confluent, confluent, early myotubes and myotubes stages. Graphs present expression of myogenic genes namely myoD, myogenin, myf6 and muscle maturation genes namely igf 1, smyhc1 and fmyhc4. Data collected from n=3 biological replicates and error bars represent standard deviation. * denotes significant difference with p < 0.05.

**Figure 4**: Graph represents frequency distribution of myotubes with number of nuclei per myotube ranging from 3 to 8+ nuclei per myotubes compared between early and late myotubes stage myotubes in monolayer. Data was generated from n=20 isolations and n=6 randomly selected images from each isolation, error bars represent the standard deviation.

**Figure 5:** Representative macroscopic images of collagen gels at two different time points embedded with zebrafish skeletal muscle cells. **A**) Collagen gel at day 1, **B**) Collagen gel after 12 days. **C**) Graph representing gel contraction or remodelling (bowing) of cellular matrix at the middle (highlighted with black arrows) due to passive force generated by zebrafish MPC’s over a time of 12 days. Data in the graph was collected from n=6 gels over the time and error bars represents standard deviation.

**Figure 6:** Representative immuno-stained zebrafish skeletal muscle cells in 3D tissue engineered collagen constructs at pre- and post- differentiation time points. Desmin protein is stained in red and nuclei in blue **A)** zebrafish MPC’s at day 1 before differentiation, **B)** zebrafish MPC’s after 5 days in differentiation. **C)** Graphs representing number of myotubes at pre- and 5 days post- differentiation per microscopic frame. **D)** Graph represents number of myotubes against number nuclei per myotube at 5 days post differentiation stage. **E)** Fusion index comparison of zebrafish MPC’s cultured in monolayer and 3D collagen constructs. Differentiated multinucleated myotubes are highlighted with the use of white arrows in image **B**. Data presented graphs was generated from n=6 collagen constructs and n=20 image per construct, error bars represent standard deviation with significance set at p < 0.05.

**Figure 7:** Relative mRNA expression graphs from 3D collagen constructs embedded with zebrafish skeletal muscle cells sampled before differentiation and myotube stage relative to after isolation time point. Graphs present expression of myogenic genes namely myoD, myogenin, myf6 and muscle maturation genes namely igf 1, smyhc1 and fmyhc4. Data collected from n=3 biological replicated each time point and error bars represent standard deviation. * denotes significant difference with p < 0.05.
