## Supplementry file 1 for "Zebrafish skeletal muscle cell cultures: Monolayer to three-dimensional tissue engineered collagen constructs"

**Supplementary file 1:** Representative images of zebrafish MPC’s embedded in 3D collagen constructs at seeding densities i.e. 2, 4, 6, 8, and 10 million cells/gel after four days. 10 million cells/gel-1 is an image of 3D collagen construct near the A-frame and 10 million cells/gel-2 is captured from the middle of the gel. All images are taken at 100X magnification.

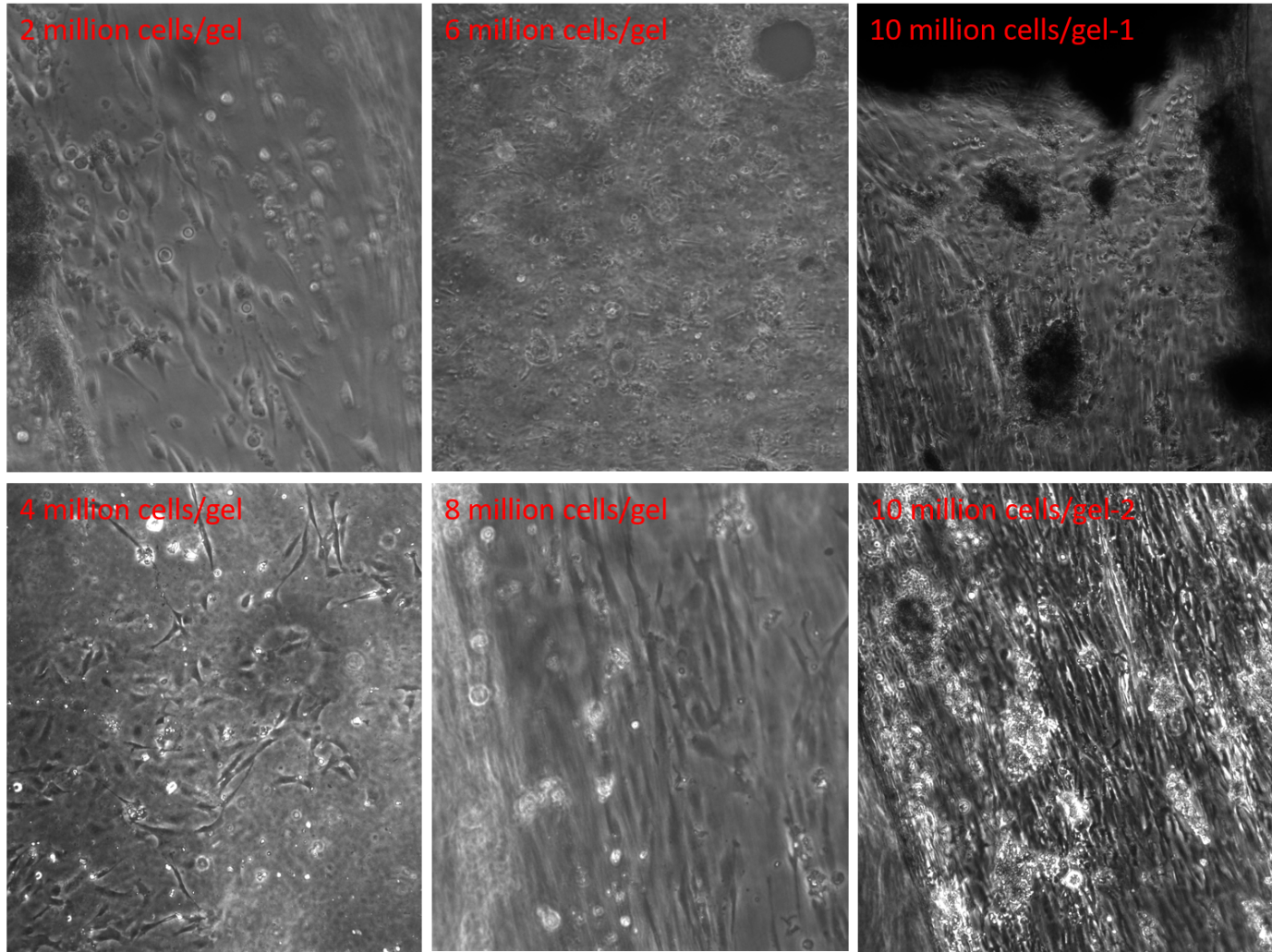

Protocol for cryo-freezing of zebrafish skeletal muscle cells:

To freeze zebrafish muscle cells, the cell pellet was obtained via centrifugation, the supernatant was removed and the cell pellet re-suspended in cryo-freezing solution constituted of 90% FBS and 10% Dimethyl sulfoxide (DMSO). After thorough mixing by pipetting, the cell suspension was transferred to a 1.8ml cryovial (Fisher Scientific) at million cells per cryovial, and placed in a “Mr Frosty” (Fisher Scientific). The “Mr Frosty” was then placed in a -80°C freezer overnight. The “Mr Frosty” containers allow for the slow (-1°C/minute) freezing of the cell suspension. The vials were then transferred to liquid nitrogen for storage. When cells were resuscitated from cryopreservation, vials were quickly thawed and plated at an appropriate density on gelatin (Sigma, UK) coated six well plates with coverslips in each well.
