## Supplementary material for "Zebrafish skeletal muscle cell cultures: Monolayer to three-dimensional tissue engineered collagen constructs": Table 1

**Table 1:** Primer sequences used for qrt-PCR

| **Target mRNA** | **Primer sequence (5’-3’)** | **Product size (in bp)** | **Annealing temp. (⁰C)** |
| --- | --- | --- | --- |
| *myoD* | F= GGCTGCCCAAAGTGGAGATTCTGA  R= TGGGCCCATAAAATCCATCATGCCA | 137 | 59 |
| *myogenin* | F= GCTCCACATACTGGGGTGTCGT  R= AGATCCTCGTGGGCGGAGCT | 125 | 59 |
| *myf6* | F= GCAGGACCTCTTGCATTCGCTGG  R= AGACTCCAACACGGCTCCTTCTC | 193 | 59 |
| *smyhc1* | F= AGAGGCTGAGGAACAGGCCA  R= CCTTTCTTGGGTCCTGAATCACGG | 147 | 57 |
| *fmyhc4* | F= GGGAGAAGGCCAAGAAGGCCA  R= CAGACGGTGCTGCAGGTCCT | 137 | 58.5 |
| *igf1* | F: GGGATGTCTAGCGGTCATTT  R: CTACATGCGATAGTTTCTGC | 489 | 58.5 |
| *ef1 α* | F= CTGGAGGCCAGCTCAAACAT  R= ATCAAGAAGAGTAGTACCGCTAGCATTAC | 87 | 60 |
| *β Actin* | F= CGAGCTGTCTTCCCATCCA  R= TCACCAACGTAGCTGTCTTTCTG | 86 | 59 |

Abbreviations

RT-PCR: Reverse transcription polymerase chain reaction; mRNA: Messenger RNA; bp: base pairs; temp.: temperature; F: forward strand; R: reverse strand.
