## Supplementary material for "Zebrafish skeletal muscle cell cultures: Monolayer to three-dimensional tissue engineered collagen constructs": Table 2

**Table 2**: Comparative morphological characteristics such as fusion index, number of myotubes per image, myotube length and width at early and late differentiation time points in monolayer. Data was generated from six randomly selected images/isolation, n=20 isolations, presented as means ± standard deviation. * denotes significant difference among the groups set at p < 0.05.

|  | **Fusion index** | **Number of Myotubes** | **Myotube length** | **Myotube width** |
| --- | --- | --- | --- | --- |
| *Early* | 60.03 ± 5.21 | 5.67 ± 1.21 | 336.87 ± 7.01 | 8.30 ± 0.21 |
| *Late* | 56.60 ± 4.92 | 5.86 ± 1.11 | 360.92 ± 7.55* | 10.80 ± 0.27* |
